# First-Trimester Non-Invasive Prediction of Preterm Birth Using Cell-Free DNA Fragmentomics

**DOI:** 10.64898/2026.07.07.736241

**Authors:** My-Diem Nguyen Pham, Minh-Tam Thi Phan, Nhat-Thang Tran, Ta-Son Vo, Hong-Thinh Le, Thu-Ha Thi Nguyen, Quoc-Huy Vu Nguyen, Minh-Thi Thi Ha, Tam Minh Le, Diem-Tuyet Thi Hoang, Khanh-Trang Nguyen Huynh, Nhan Viet Nguyen, Chuong Canh Nguyen, Thuong Chi Bui, Xuan Thanh Nguyen, Sa Viet Le, Vinh Dinh Tran, My-Nhi Ba Nguyen, Thong Van Nguyen, Tuyet-Anh Thi Nguyen, Ba Phuoc Hoang, Trong Van Nguyen, Thuy-Ai Thuy Nguyen, Toa Tri Nguyen, Thang Duc Duong, Cuong Huy Pham, Kim-Oanh Thi Luong, Cuong Ngoc Dao, Khanh Van Hoang, Thu-Thanh Thi Huynh, Khuong Manh Nguyen, Son-Tra Thi Tran, Hoanh Trung Tran, Son Canh Nguyen, Thuy Dinh Tran, Phương Thi Lan Nguyen, Thanh Viet Pham, Kong Chi Pham, Minh Doan Thai, Thanh-Thuy Thi Do, Hoa Thi Dao, Yen-Linh Thi Van, Sang Hung Tang, Hoai-Nghia Nguyen, Minh-Duy Phan, Hoa Giang, Cuong Thach Nguyen, Quang Thinh Trac

## Abstract

**Objective:** To develop and validate a cell-free DNA (cfDNA) fragmentomic classifier for the early prediction of spontaneous preterm birth (PTB) using routine first-trimester non-invasive prenatal testing (NIPT) data.

**Methods:** A nested case-control study was conducted within a prospective multicenter Vietnamese cohort comprising 286 pregnancies, including 82 spontaneous PTB cases and 204 term controls. Maternal plasma cfDNA collected during routine first-trimester NIPT (median gestational age, 12 weeks) was sequenced to a depth of approximately 20 million reads per sample. Five fragmentomic feature categories including copy number alterations, end-motif composition, nucleosome distance, fragment length, and joint fragment-length×end-motif were evaluated for PTB prediction. Machine learning classifiers were developed in a training cohort (n = 228, 65 PTB vs 163TB) and tested in a validation cohort (n = 58, 17 PTB vs 41 TB).

**Results:** Among the five fragmentomic feature classes evaluated, 4-mer end-motif (EM) profiles exhibited the most pronounced differences between PTB and term control samples. Consistent with these findings, the EM-based classifier demonstrated the highest discriminative performance in the validation cohort, achieving an AUC of 0.970 (95% CI, 0.912–1.000). At a specificity >90%, the model achieved a sensitivity of 94% (95% CI, 78–100%).

**Conclusion:** These findings demonstrate that cfDNA EM signatures derived from routine first-trimester NIPT can accurately identify pregnancies at risk of spontaneous preterm birth, without additional blood collection or sequencing, thereby extending the clinical utility of existing prenatal screening infrastructure.

**KEY POINTS:** *What is already known about this topic?:*

- Current first-trimester prediction strategies based on maternal characteristics, cervical length, and biochemical markers have limited predictive accuracy, particularly in nulliparous women.
- Existing cfDNA-based approaches have shown only modest performance or require additional assays, limiting clinical applicability.

*What does this study add?:*

- Existing NIPT sequencing data can be repurposed (without additional blood sampling or sequencing) for accurate prediction of spontaneous preterm birth (AUC=0.970).
- A classifier employing 4-mer end-motif (EM) profiles achieved an AUC of 0.970. At a specificity >90%, the model achieved a sensitivity of 94%.

## INTRODUCTION

Preterm birth (PTB), defined as delivery before 37 completed weeks of gestation, is a leading cause of neonatal morbidity and mortality worldwide, affecting ∼11% of live births and accounting for 35% of pregnancy-related deaths. In 2020, an estimated 13.4 million infants were born preterm (9.9% of live births), with no measurable global decline over the previous decade^1^. The burden is concentrated in Southern Asia and sub-Saharan Africa, which together account for ∼65% of cases and have the most limited capacity for early risk identification^1^. Most PTBs result from spontaneous preterm labor with intact membranes or preterm premature rupture of membranes. PTB is a heterogeneous obstetric syndrome driven by multiple convergent pathways, including intra-amniotic inflammation, decidual senescence, vascular dysfunction, breakdown of maternal-fetal immune tolerance, and premature activation of the parturition cascade^2,3^. Despite decades of intensive research, accurate prediction of spontaneous preterm birth (PTB) in asymptomatic women during early gestation remains a critical unmet clinical need. Contemporary risk stratification relies predominantly on maternal obstetric history, sonographic cervical length measurement, and select biochemical markers, including fetal fibronectin and the sFlt-1/PlGF ratio^4^. However, these approaches exhibit only modest sensitivity and specificity, particularly in the first and early second trimesters, and fail to identify a substantial proportion of nulliparous women who subsequently deliver preterm^4^.

Maternal plasma cell-free DNA (cfDNA), released mainly through apoptosis of placental trophoblasts and maternal hematopoietic cells, serves as a dynamic real-time sensor of the maternal-fetal interface^5^. The rapidly advancing field of cfDNA fragmentomics has revealed highly non-random fragmentation patterns (copy number alterations, nucleosome distance, fragment length) in circulating cfDNA, most notably the characteristic 4-mer nucleotide signatures at fragment ends (end motifs, EMs), along with distinct pathophysiological footprints that are detectable across diverse pregnancy complications^6,7^. Existing cfDNA-based approaches to PTB prediction have required additional assays beyond routine NIPT. A transformer-based architecture integrating cfDNA with cfRNA achieved an external AUC of 0.890, but required paired cfRNA sequencing that substantially increases per-sample assay cost^8^. Elevated cfDNA fetal fraction in the second trimester has shown only modest and inconsistent predictive performance^9,10^, and a recently reported single-layer promoter-profiling classifier achieved an external AUC of 0.849 using samples obtained between 12 and 28 weeks of gestation^11^. Most recently, a multimodal classifier integrating cfDNA fragmentomic and epigenetic features achieved good predictive performance for preterm preeclampsia, with an AUROC of 0.85^12^. In this proof-of-concept study, we evaluated whether fragmentomic features derived from routine first-trimester NIPT cfDNA can discriminate pregnancies at risk for spontaneous PTB from uncomplicated term pregnancies, in a multi-center Vietnamese cohort.

## METHODS

### Participant Recruitment

This prospective multicentre study included 286 pregnant women recruited from multiple clinical sites across Vietnam who met the eligibility criteria and completed the study protocol. All participants underwent routine first-trimester non-invasive prenatal testing (NIPT) and were enrolled before the onset of clinical signs of preterm labour. Participants were prospectively followed until delivery, and pregnancy outcomes were systematically recorded. Peripheral blood samples were collected at the time of routine first-trimester NIPT, before the onset or diagnosis of spontaneous preterm birth.

By definition, term controls delivered at ≥ 37 completed weeks of gestation while spontaneous PTB was defined as delivery before 37 weeks of gestation associated with cervical dilation and/or premature rupture of membranes. Pregnancies resulting in medically indicated preterm delivery, including those related to preeclampsia, fetal growth restriction, or maternal medical conditions, were excluded to focus specifically on spontaneous PTB

### Sample Collection and cfDNA Extraction

Maternal peripheral blood was collected into cell-free DNA blood collection tubes (BD, USA). Plasma was isolated by a two-step centrifugation protocol consisting of an initial centrifugation at 2,000 × g for 10 min at 4°C, followed by a second centrifugation at 16,000 × g for 10 min at 4°C to remove residual cellular debris. The resulting cell-free plasma was aliquoted into barcoded tubes and stored at −80°C until analysis.

Cell-free DNA (cfDNA) was extracted using the MagMAX™ Cell-Free DNA Isolation Kit (Thermo Fisher Scientific, USA) according to the manufacturer’s instructions. cfDNA concentration was measured using the QuantiFluor® dsDNA System and Quantus™ Fluorometer (Promega, USA). Purified cfDNA was eluted in 30 μL of elution buffer and stored at −20°C until library preparation. Samples with cfDNA concentrations ≥0.05 ng/μL were considered to have passed quality control and were included in downstream analyses. Assessment of the A260/A280 ratio was not performed because cfDNA concentrations were below the optimal range for reliable NanoDrop spectrophotometric measurement.

### Whole-genome Sequencing of plasma cfDNA

Plasma cfDNA libraries were prepared using the NEBNext® Ultra™ II DNA Library Prep Kit for Illumina (New England Biolabs, USA), followed by AMPure XP bead-based purification and size selection (Beckman Coulter, USA). Libraries were normalized, pooled, and sequenced on the AVITI™ platform (Element Biosciences, USA) to a depth of approximately 20 million paired-end reads (2 × 75 bp) per sample.

*Batch-effect control*. To minimise technical confounding of fragmentomic features, spontaneous PTB cases and term controls were processed together throughout the workflow rather than in outcome-defined batches. Plasma extraction, library preparation, and sequencing were performed by operators blinded to pregnancy outcome. Since maternal plasma was collected during routine first-trimester NIPT, well before pregnancy outcome was known, sample processing was inherently independent of case–control status. All libraries were prepared with the identical chemistry and sequenced on the same AVITI platform to a uniform target depth of approximately 20 million paired-end reads (2 × 75 bp) per sample, and all samples were processed through a single bioinformatic pipeline.

### Sequencing data preprocessing

Raw FASTQ files underwent quality control with FastQC (v0.12.1). Adapters and low-quality bases were trimmed using Trimmomatic (v0.39)^13^. The filtered reads were then aligned to the GRCh38/hg38 human reference genome using BWA-MEM2 (v2.3). Duplicate reads were marked and removed with Picard MarkDuplicates (v4.0.1)^14^. Fragmentomic features were subsequently extracted from the deduplicated, aligned BAM files.

### Fragmentomic feature extraction

Fragment end motifs (EMs) were defined as the 4-mer genomic sequences immediately adjacent to the 5′ end of aligned fragments. Relative frequencies of all 256 possible 4-mers were calculated and normalized to the total fragment count, yielding a 256-dimensional feature vector per sample. These vectors constituted the primary EM feature set for downstream model development.

Nucleosome distance (ND) features were extracted using the Griffin reference panel^15^. Copy number alteration (CNA) features were derived from read-depth analysis across genome-wide 1,000-bp bins. Fragment length (FLEN) features were generated from the size distribution of aligned cfDNA fragments.

### Model development and validation

The prediction model was constructed as an L1-regularized (LASSO) logistic regression classifier using scikit-learn (v1.9.0) to distinguish spontaneous preterm birth (PTB) from control (CTRL) pregnancies. The model was trained on a development cohort of 65 PTB and 163 CTRL samples, with hyperparameter optimization including L1 regularization strength performed via 5-fold cross-validation. The final model was then independently evaluated, without retraining, on a validation cohort comprising 17 PTB and 41 CTRL samples. For direct comparison, models based on fragmentomic feature sets were trained and assessed using the identical pipeline.

Model performance was evaluated using the area under the receiver operating characteristic curve (AUC-ROC) in a validation cohort. To reflect a clinically relevant screening threshold with high specificity, sensitivity and specificity were additionally reported at a fixed specificity of > 90%, prioritizing low false-positive rates suitable for population-level screening.

## RESULTS

### Cohort characteristics and study design

To develop a non-invasive approach for early prediction of spontaneous PTB, we leveraged routine NIPT data from a prospective multicentre nested case–control study conducted across independent Vietnamese clinical sites. Following pregnancy outcome ascertainment, 286 eligible pregnancies were included for cfDNA profiling, comprising 82 spontaneous PTB cases (28.7%) and 204 uncomplicated term pregnancies (71.3%) (**Figure 1A**). Maternal plasma samples were collected during routine first-trimester NIPT screening at a median gestational age of 12 weeks, thereby establishing a predictive window well in advance of clinically apparent preterm labour. To support model development and evaluation of performance, the cohort was stratified into two mutually exclusive datasets: a training cohort (n = 228; 65 PTB and 163 TB) a validation cohort (n = 58; 17 PTB and 41 TB) (**Figure 1A**).

**Figure 1.**
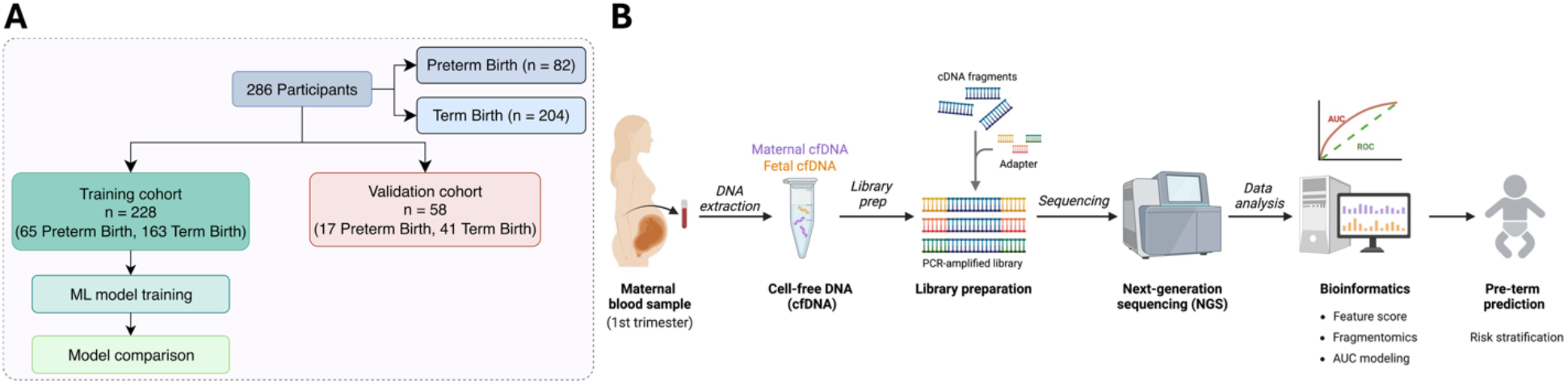
Workflow and Study Design. **A**) A total of 286 participants, including 82 spontaneous PTB cases and 204 term birth (TB) controls, were randomly assigned to a training cohort (n = 228; 65 PTB and 163 TB) for machine learning model development and comparative performance evaluation, and a validation cohort (n = 58; 17 PTB and 41 TB) for assessment of model generalizability. **B**) Maternal plasma samples collected during routine first-trimester NIPT were sequenced to a depth of approximately 20 million reads per sample on the AVITI platform and analyzed using a multi-layer cfDNA fragmentomic workflow incorporating end-motif composition, fragment length, nucleosome positioning, and copy number alterations features for PTB risk prediction.

Plasma-derived cfDNA underwent whole-genome sequencing on the AVITI platform, generating approximately 20 million sequencing reads per sample (**Figure 1B**). Baseline demographic and obstetric characteristics were broadly balanced across the training and validation cohorts (**Table 1**). No significant differences were observed between spontaneous PTB and term birth (TB) groups with respect to gestational age at blood collection, maternal age, body mass index, height, or parity (*P* > 0.05). The study population was ethnically homogeneous, with all participants self-identifying as Kinh. To capture diverse dimensions of cfDNA fragmentation biology, we extracted five classes of fragmentomic features from each sequencing dataset: fragment length distribution (50–300 bp), nucleosome distance profile (−300 to +300 bp), CNA profile, 4-mer end-motif representation (256 motifs), and joint fragment-length ×end-motif (FLE ×*EM*). These feature modalities were subsequently evaluated as predictive inputs for machine-learning models aimed at differentiating spontaneous PTB from term births.

**Table 1:** Cohort characteristics.

| Characteristic | Training cohort |  |  | Validation cohort |  |  |
| --- | --- | --- | --- | --- | --- | --- |
|  | PTB | TB | Difference (P value) | PTB | TB | Difference (P value) |
| N | 65 | 163 |  | 17 | 41 |  |
| Ethnic | Kinh | Kinh |  | Kinh | Kinh |  |
| Age (years mean ± s.d.) | 31.4 ± 5.4 | 31.6 ± 4.0 | 0.801 | 30.3 ± 2.5 | 32.0 ± 3.1 | 0.139 |
| Multi-fetal pregnancy | 1 (1.5%) | 0 (0.0%) | 0.285 | 0 (0.0%) | 0 (0.0%) | 1.000 |
| Height (centimeters mean ± s.d.) | 156.5 ± 4.9 | 156.5 ± 5.1 | 0.907 | 155.2 ± 5.1 | 156.2 ± 4.4 | 0.476 |
| BMI (mean ± s.d.) | 22.5 ± 3.7 | 21.7 ± 3.6 | 0.080 | 22.0 ± 4.9 | 21.7 ± 3.6 | 0.755 |
| Gravidity (mean ± s.d.) | 1.8 ± 1.0 | 2.1 ± 1.1 | 0.139 | 1.6 ± 0.8 | 2.0 ± 1.1 | 0.226 |
| Nulliparous (N (%)) | 22 (33.8%) | 53 (32.5%) | 0.509 | 7 (41.2%) | 13 (31.7%) | 0.311 |
| PTB history (N (%)) | 0 (0.0%) | 0 (0.0%) | 1.000 | 1 (5.9%) | 0 (0.0%) | 1.000 |
| Abortion history (N (%)) | 9 (13.8%) | 36 (22.1%) | 0.258 | 0 (0.0%) | 8 (19.5%) | 0.174 |
| Threatened PTB (N (%)) | 34 (51.5%) | 0 (0.0%) | <0.001 | 11 (64.7%) | 0 (0.0%) | <0.001 |
| GA at sampling (median weeks (range)) | 12.0 (7.0–13.1) | 12.0 (6.0–16.1) | 0.064 | 12.0 (6.9–13.0) | 12.0 (6.7–13.0) | 0.836 |
| IVF (N (%)) | 3 (4.6%) | 20 (12.3%) | 0.093 | 0 (0.0%) | 5 (12.2%) | 0.308 |
| Cesarean (N (%)) | 0 (0.0%) | 5 (3.1%) | 0.325 | 0 (0.0%) | 1 (2.4%) | 1.000 |

### Genome-wide fragmentomic features distinguish PTB from term controls

To characterise the genome-wide fragmentomic alterations underlying first-trimester PTB, we compared end-motif composition, fragment-size distribution, nucleosome positioning, FLEN × EM, and copy-number profiles between PTB and term control (CTRL) samples (**Figure 2**). The genome-wide 4-mer end-motif landscape exhibited coordinated shifts in motif frequency between groups **(Figure 2A**): G*** and T*** motifs were broadly enriched in CTRL but depleted in PTB, whereas C*** motifs showed reciprocal enrichment in PTB; A*** motifs exhibited more variable distributions without a consistent disease-associated pattern. At the individual motif level (**Figure 2B**), the five motifs most significantly increased in PTB were CCAA, CAAA, CTAA, CAAC, and AAAC, while the five most significantly decreased were TTGG, TCGG, TCGC, GCGG, and TCGT (all *P* < 0.05, Wilcoxon rank-sum test); CCAA and TTGG showed the largest positive and negative differences, respectively.

**Figure 2.**
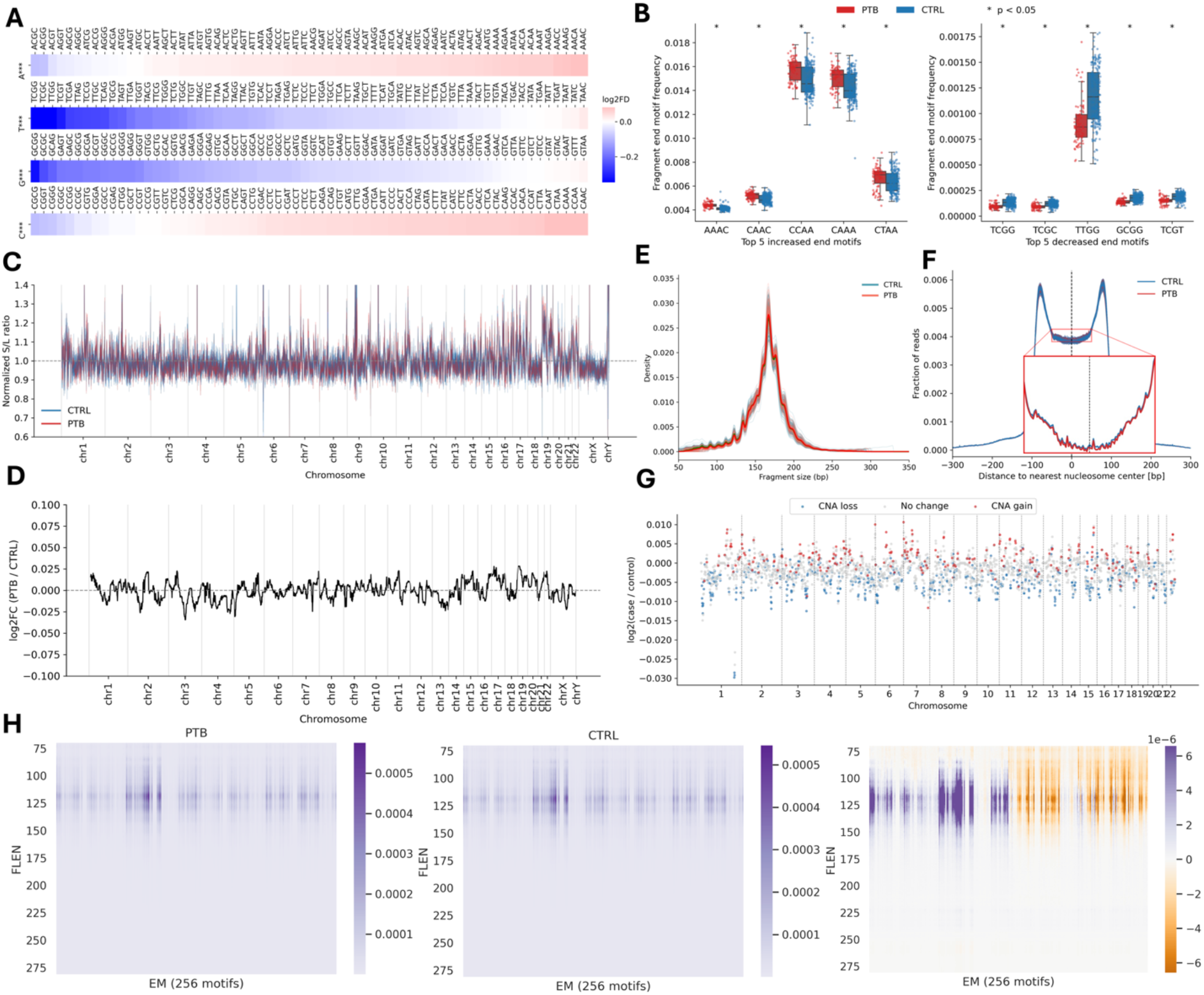
Fragmentomic Features in Preterm Birth and Term Controls. **A**) Genome-wide end-motif (EM) profiles showed marked 4-mer composition shifts between PTB and CTRL, with G***/T*** motifs enriched in CTRL and C*** motifs enriched in PTB. **B**) Differential 4-mer motifs distinguished PTB and CTRL, with CCAA/CTAA enriched and TCGG/TCGT depleted in PTB (P < 0.05). **C**) Short-to-long fragment ratios showed no consistent group-specific chromosomal differences. **D**) Nucleosome-depleted region (NDR) profiles remained centered around zero, indicating similar chromatin accessibility. **E**) Fragment length profiles showed a conserved ∼167 bp peak, with increased sub-nucleosomal fragments in PTB. **F**) Inter-nucleosome distance distributions were similar between groups. **G**) CNA profiles showed no consistent differences between PTB and CTRL. **H)** Joint fragment-length ×end-motif (FLEN×EM) profiles for PTB, CTRL, and their difference (PTB − CTRL) showed motif-specific enrichment concentrated in the 100–150 bp sub-nucleosomal fragment range, with certain 4-mer motifs more depleted (purple, left/middle panels) or differentially enriched (orange, right panel) in PTB relative to CTRL.

Fragment-size and coverage-based features show only modest differences. Genome-wide normalised short-to-long (S/L) fragment ratios oscillated around the reference baseline in both groups without consistent chromosome-specific alterations (**Figure 2C**), and genome-wide log_2_(PTB/CTRL) nucleosome-depleted region (NDR) profiles remained centred around zero (range: −0.10 to +0.10), indicating broadly conserved chromatin accessibility (**Figure 2D**). The cfDNA fragment-size distribution showed the characteristic mono-nucleosomal peak at ∼167 bp in both groups, with PTB samples exhibiting a higher peak density and a modest enrichment of sub-nucleosomal fragments (100–145 bp) (**Figure 2E**), and inter-nucleosome distance profiles displayed the expected bimodal pattern at approximately ±100 bp from the nearest nucleosome dyad without inter-group divergence (**Figure 2F**). Genome-wide CNA analysis revealed randomly distributed gain and loss events across all chromosomes in both groups, without consistent disease-associated patterns (**Figure 2G**), supporting the conclusion that the discriminative cfDNA signal for first-trimester PTB does not reside in chromosomal copy-number alterations. In addition, joint two-dimensional (2D) analysis of FLEN × EM revealed that the PTB-associated end-motif alterations were spatially concentrated within specific fragment-size compartments (**Figure 2H**). Both PTB and CTRL samples displayed the expected dominant end-motif signal in the 160–180 bp mono-nucleosomal fragment range; however, the differential (PTB vsv CTRL) profile demonstrated that the largest inter-group divergence was localised to the 100–150 bp sub-nucleosomal fragment range, with a subset of 4-mer motifs showing pronounced enrichment (orange) and others pronounced depletion (purple) in PTB relative to CTRL.

### Comparative performance of fragmentomic classifiers for preterm birth prediction

To identify the fragmentomic feature class with the strongest discriminative power for first-trimester spontaneous PTB, we benchmarked five classifiers built on EM, CNA, ND, FLEN, and combined FLEN × EM features through 5-fold cross-validation in the training cohort, followed by evaluation in a validation cohort (**Figure 3**). In the training cohort, end motifs achieved the highest cross-validated discriminative performance. Specifically, the EM-based classifier attained a median per-fold AUC of 0.980, substantially exceeding FLEN (0.712), CNA (0.698), FLEN × EM (0.648), and ND (0.587) (**Figure 3A**). At a fixed specificity threshold of ≥ 90%, EM achieved a sensitivity of 0.94 with specificity of 0.96, while the other length- and coverage-based features showed clinically inadequate sensitivities despite preserved specificities (CNA: 0.25/0.95; ND: 0.12/0.97; FLEN: 0.29/0.94; FLEN × EM: 0.22/0.95) (**Figure 3B**).

**Figure 3.**
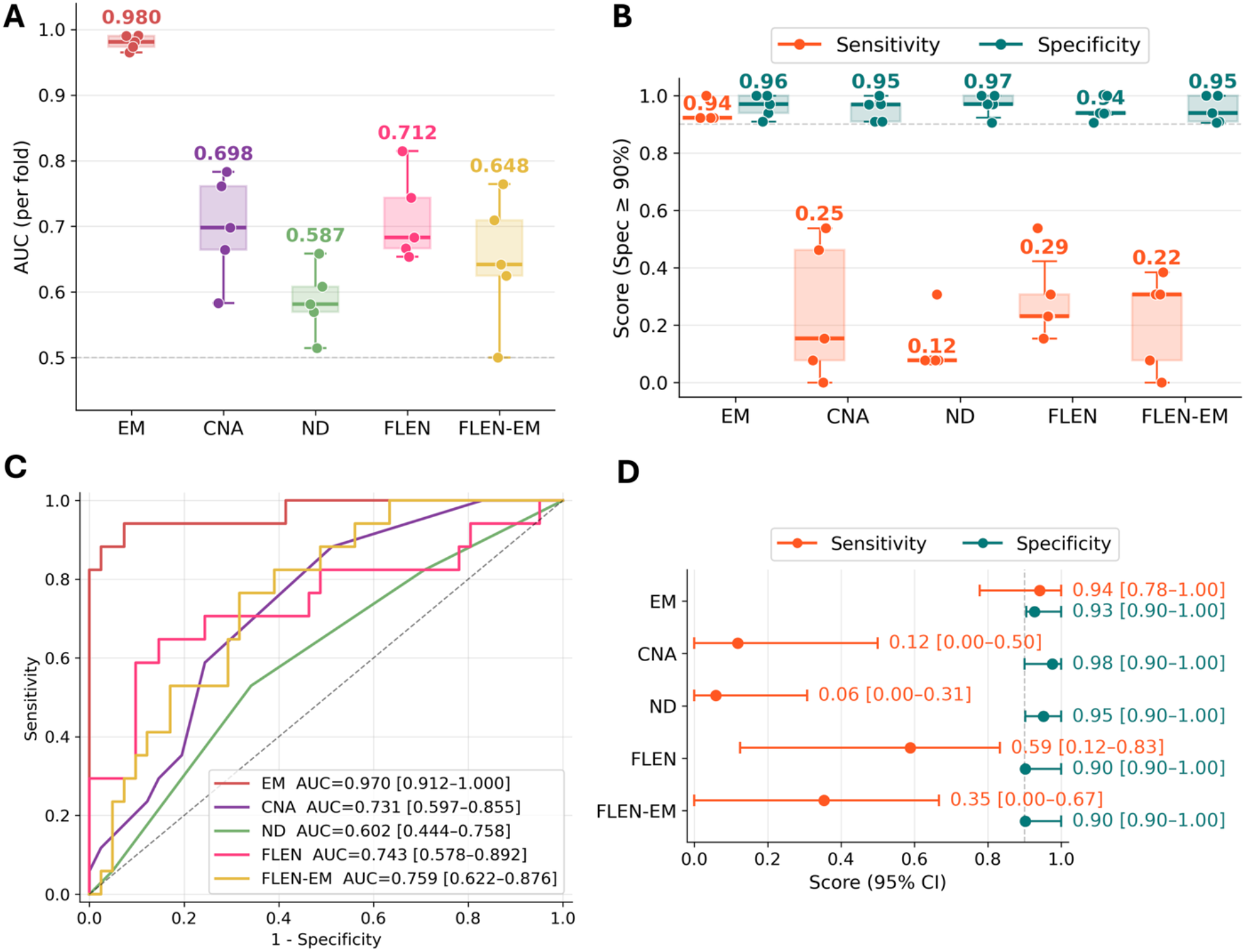
Comparative performance of fragmentomic classifiers for preterm birth prediction. **A)** AUC distribution per cross-validation fold in the training cohort for five classifiers built on end motif (EM), copy-number alteration (CNA), nucleosome distance (ND), fragment length (FLEN), and combined FLEN-EM features. Median AUC values are annotated above each box. **B**) Sensitivity (orange) and specificity (teal) per cross-validation fold in the training cohort at a fixed specificity threshold of ≥ 90%. **C**) Receiver operating characteristic (ROC) curves for the five classifiers evaluated in the validation cohort. Area under the curve (AUC) with 95% confidence intervals is shown for each feature. Diagonal dashed line indicates the line of no discrimination. **D**) Sensitivity and specificity with 95% confidence intervals in the validation cohort at a fixed specificity threshold (dashed line) of ≥ 90%.

Consistently, EM-based discrimination was preserved in the validation cohort. The EM classifier achieved a test-set AUC of 0.970 (95% CI: 0.912–1.000), with a sensitivity of 0.94 (0.78–1.00) and specificity of 0.93 (0.90–1.00). In contrast, other features showed substantially lower discriminative performance (CNA: AUC 0.731 [0.597–0.855], sensitivity 0.12 [0.00–0.50]; ND: AUC 0.602 [0.444–0.758], sensitivity 0.06 [0.00–0.31]; FLEN: AUC 0.743 [0.578–0.892], sensitivity 0.59 [0.12–0.83]). Notably, the combined FLEN × EM model (AUC 0.759 [0.622– 0.876]; sensitivity 0.35 [0.00–0.67]) underperformed the EM-only classifier, suggesting that integration of low-discriminative FLEN features introduced noise that degraded the high-quality EM signal rather than providing complementary information (**Figure 3C-D**). The reproducibility of EM performance between cross-validated training (AUC 0.980) and validation cohorts (AUC 0.970), together with the substantial performance gap between EM (AUC > 0.97) and other length- or coverage-based features (AUC range 0.60–0.76), establishes EM as the dominant fragmentomic discriminator for first-trimester PTB.

## DISCUSSION

In this prospective multicentre nested case-control study of 286 pregnant women enrolled across multiple clinical sites in Vietnam, we demonstrated that a cfDNA fragmentomic approach applied to routine first-trimester NIPT samples achieves highly accurate prediction of spontaneous preterm birth. Among five candidate fragmentomic feature sets evaluated, including end motif (EM), fragment length (FLEN), copy-number alteration (CNA), nucleosome distance (ND), and the combined FLEN × EM, the EM-based classifier achieved the highest discriminative performance, with an AUC of 0.970 (95% CI: 0.912–1.000) in the validation cohort. At a fixed specificity threshold of ≥ 90%, the EM classifier maintained a sensitivity of 0.94 (95% CI: 0.78–1.00), substantially exceeding the sensitivities achieved by other length- and coverage-based features (FLEN 0.59, CNA 0.12, ND 0.06, FLEN × EM 0.35).

The performance advantage of EM over other length- and coverage-based features is biologically interpretable. The 4-mer composition at cfDNA fragment termini directly reflects the cleavage preferences of extracellular endonucleases including DNASE1, DNASE1L3, and DFFB^16^ and therefore encodes a biochemical signature of the systemic and placental^16^ processes generating circulating cfDNA, whereas fragment length, copy-number, and nucleosome positioning aggregate signal across the genome and are less sensitive to focal maternal–fetal perturbations^6,7,17^. The substantial performance gap in our cohort (EM AUC 0.970 versus FLEN 0.743, CNA 0.731, ND 0.602, FLEN × EM 0.759) supports end-motif composition as the primary fragmentomic discriminator of first-trimester PTB and indicates that integration of low-discriminative features introduces noise rather than complementary signal.

In the most comparable prior study, PTerm, a promoter-profiling classifier developed in a large Chinese cohort, reported an AUC of 0.849 in external validation using samples collected between 12 and 28 weeks of gestation^11^. A subsequent transformer-based model integrating cfDNA and cfRNA achieved an external AUC of 0.890; however, it required additional PALM-Seq cfRNA sequencing, which substantially increases cost and is incompatible with standard NIPT workflows^8^. Earlier studies relying on cfDNA fetal fraction showed only modest and inconsistent performance^9,18,19^. Furthermore, the DREAM challenge benchmarking of maternal multi-omics data demonstrated that no single analyte class provides robust early-pregnancy PTB prediction, with AUROCs ranging from 0.60 to 0.76 for samples collected at 27–33 weeks^20^. In the most recent study, a multimodal cfDNA classifier integrating fragmentomic and epigenetic features achieved an AUROC of 0.85 for preterm preeclampsia^12^. In our study, the EM-based classifier developed here achieved markedly superior discriminative performance (AUC 0.970, sensitivity 0.94 at ≥90% specificity) using only the cfDNA data already generated by routine first-trimester NIPT, representing a substantive improvement in both predictive accuracy and translational feasibility.

From a clinical standpoint, the EM-based classifier offers substantial translational potential due to its seamless integration with the established NIPT infrastructure, which performs over 10 million tests annually in more than 60 countries^21^. By utilizing the existing cfDNA sequencing data generated for routine fetal aneuploidy screening, the model requires no additional blood sampling, library preparation, or sequencing. These attributes establish a new standard for feasible implementation in routine antenatal care, particularly in low- and middle-income countries that bear the greatest burden of preterm birth yet possess the most constrained biomarker resources^1^. Moreover, reliable first-trimester prediction creates a critical window for timely preventive interventions, such as vaginal progesterone supplementation, cervical pessary placement, and enhanced antenatal monitoring in high-risk pregnancies.

Several limitations warrant explicit acknowledgement. First, the cohort comprised exclusively Vietnamese (Kinh ethnicity) participants enrolled across multiple sites within a single healthcare system; generalisability to other ancestries and healthcare settings requires prospective external validation in geographically and ethnically diverse cohorts. Second, although the EM-based classifier was evaluated in a validation cohort, the modest overall cohort size (n = 286) relative to the 256-dimensional EM feature space raises the possibility of optimistic performance estimation, and the high AUC obtained (0.970, 95% CI 0.912–1.000) should be interpreted with appropriate caution pending replication in larger independent cohorts. No formal sample size or power calculation was conducted, consistent with the exploratory nature of this study. Larger prospective studies will be essential to externally validate the performance of the classifier, evaluate its generalizability across populations, assess robustness against potential confounders.

In conclusion, this proof-of-concept study demonstrates that end-motif fragmentomic features derived from routine first-trimester NIPT cfDNA enable highly accurate prediction of spontaneous preterm birth, substantially outperforming length- and coverage-based features in a multicentre Vietnamese cohort. Prospective multi-ancestry validation in larger independent cohorts, combined with integration with established clinical predictors such as obstetric history and cervical length measurement, will be required before clinical implementation. The analytical framework introduced here is, in principle, extensible to other pregnancy complications with placental and immune-mediated pathophysiology, including preeclampsia, gestational diabetes, and fetal growth restriction.

## ACKNOWLEDGEMENTS

Not Applicable.

## ETHICS STATEMENT

Ethical approval for this study was obtained from the Institutional Review Board of the Medical Genetics Institute (Decision No. 3/2024/QÐ-VDTYH) and the Ethics Committee of the University of Medicine and Pharmacy at Ho Chi Minh City (Approval No. 525/HÐÐÐ-ÐHYD). Written informed consent was obtained from all participants before study enrollment, and the study was conducted in accordance with the Declaration of Helsinki.

## CONSENT

Written informed consent was obtained from all participants prior to study enrollment.

## DATA AVAILABILITY STATEMENT

The source code for fragmentomic feature extraction and classifier development are available at on request. Raw sequencing data cannot be publicly shared because of participant privacy and ethical restrictions. Access to de-identified data may be granted upon reasonable request to the corresponding author, subject to review and approval by the Institutional Ethics Review Board of the Medical Genetics Institute and applicable data-sharing regulations.

